# Persistent effects of acute trauma on Pavlovian-to-instrumental transfer

**DOI:** 10.1101/2022.08.05.502959

**Authors:** Rifka C. Derman, K. Matthew Lattal

## Abstract

In humans, an acutely traumatic experience can lead to post-traumatic stress disorder (PTSD), which is often characterized by changes in anxiety and motivation months after trauma. There are few demonstrations of the persistent motivational effects of an acute stressor in rodent approaches to PTSD. In two experiments, we evaluated the persistent effects of a battery of footshocks in one context on appetitive Pavlovian conditioning, instrumental learning, and Pavlovian-to-instrumental transfer (PIT) in a different context. A battery of footshocks before appetitive training caused deficits in single-outcome PIT in male Long Evans rats. The same battery of footshocks after appetitive training, but before testing had little effect on single-outcome PIT, but there were some qualitative deficits. Although males showed more generalized fear from the aversive to the appetitive context compared to females, there were no effects of shock on sensory-specific PIT in male or female rats. In general females showed less evidence for sensory-specific PIT compared to males, who showed robust sensory-specific PIT, with clear extinction and spontaneous recovery of the sensory-specific PIT effect across test sessions. These findings show that (a) an acute trauma can have persistent effects on general motivational processes and (b) sensory-specific PIT is a useful approach for exploring sex differences in strategies for instrumental learning. We discuss implications for current approaches to stress and motivation in preclinical and clinical studies.

## Introduction

Intensely stressful events can cause long-term aberrations in learning, memory, and motivation. In humans, the experience of either intense acute or chronic stress can lead to post-traumatic stress disorder (PTSD) which is characterized by persistent intrusive memories of the trauma, avoidance of reminders of the trauma, negative changes in cognition and mood, and alterations in arousal/stress reactivity, all which persist 30 days beyond trauma (American Psychiatric Association 2013). A major difficulty with treating PTSD is that it is highly comorbid with persistent aberrations in reward-related processes. Individuals with PTSD are at greater risk of developing alcohol use, substance use, gambling, and eating disorders, and also show a higher risk of obesity (Kessler, Chiu et al. 2005, Brewerton 2007, Swinbourne and Touyz 2007, Chwastiak, Rosenheck et al. 2011, Coughlin, Kang et al. 2011, Grubbs, Chapman et al. 2019). Although there are many preclinical demonstrations of the persistent effects of chronic stressors on reward processing (Willner, Towell et al. 1987, Piazza, Deminiere et al. 1990, Ghiglieri, Gambarana et al. 1997), the effects of acute stressors has largely been studied using the stress-induced reinstatement approach, in which a stressor is applied after extinction and the effects on instrumental responding are assessed during that period of acute stress. Remarkably little is known about the effects of acute stress on reward that persist long after that stressful experience.

One approach to evaluating persistent effects of acute stress is stress-enhanced fear learning (SEFL), which is based on years of work on the cellular, molecular, and behavioral mechanisms of fear conditioning. This approach results in a persistent hyperresponsivity to a mild stressor in one context following an exposure to a battery of intense footshocks in a different context (Rau, DeCola et al. 2005, Rau and Fanselow 2009). Chronic stressors (such as maternal deprivation or chronic variable stress) often involve exposure to stressors over multiple days in multiple contexts without an easily measured memory component. SEFL consists of a temporally and contextually isolated stressful experience that has a well characterized behavioral response (freezing behavior) associated with it. Thus, the effects of memory for trauma can be evaluated and potentially dissociated from the effects of the stress of the trauma.

SEFL has been adapted to study the persistent alterations of trauma on appetitive processes by following the shock battery session with appetitive training and testing in a novel context (Meyer, Long et al. 2013, Pizzimenti, Navis et al. 2017). Meyer et al. (2013) found that the battery of footshocks led to a persistent change in alcohol drinking and Pizzimenti et al. (2017) found that the same battery of shocks promoted expression of cocaine-induced conditional place preference in mice and methamphetamine-induced reinstatement in rats. In both of these studies, a single acute stressor resulted in persistent changes in drug-seeking or -taking. However, little is known about how the persistent effects of acute trauma on appetitive learning and motivation in the realm of natural rewards. This is an important line of inquiry because it will provide valuable insights as to unique alterations to the psychological and neurobiological processes imparted by trauma alone without the interference on drug-related plasticity in these same processes.

The Pavlovian-to-instrumental transfer (PIT) procedure allows the influence of Pavlovian stimuli on instrumental behaviors to be assessed. This approach may capture aspects of general affective motivation for reward, as well as sensory-specific memories involving associations among stimuli, responses, and predicted outcomes (Cartoni, Balleine et al. 2016). In single-outcome PIT, animals are conditioned to associate a Pavlovian conditional stimulus (CS+) with an unconditional stimulus (US; or outcome, O). Separately animals are trained to associate an instrumental response (R1) with that same outcome. During testing, SO-PIT is captured when presentation of the CS+ augments instrumental responding on R1. SO-PIT is thought to rely on a general affective motivational process and depends on psychological and neurobiological processes that are distinct from another form of PIT, sensory-specific (SS) PIT. With SS-PIT animals are conditioned to form two specific Pavlovian associations (CS1-O1 and CS2-O2) and separately trained to associate two different instrumental responses with these same outcomes (R1-O1 and R2-O2). In testing, both manipulanda are available and the Pavlovian CSs are presented to assess their influence on instrumental responding. SS-PIT occurs when a given CS preferentially invigorates instrumental responding on the manipulandum previously reinforced with the same outcome compared to the manipulandum previously reinforced with the alternate outcome (i.e., S1 increases R1 more than R2 and S2 increases R2 more than R1).

Because PIT separates Pavlovian and instrumental memory processes, it is a powerful tool for investigating how trauma alters specific memory and motivational processes. Much of the work on stress and reward (such as the work on cue-induced reinstatement) cannot distinguish between Pavlovian and instrumental processes. In humans, high anxiety levels (not experimentally induced) have been associated with deficits in SO-PIT (Quail, Morris et al. 2017), whereas experimentally induced stress just prior to testing has been shown to promote SO-PIT (Pool, Brosch et al. 2015) and to not alter SS-PIT (Steins-Loeber, Lörsch et al. 2020). In rodents, acute stress just prior to testing has been shown not to affect SO-PIT expression (Pielock, Braun et al. 2013), but to transiently block SS-PIT expression (Morgado, Silva et al. 2012). In the following experiments we evaluate the persistent (>21 days) effects of acute trauma (exposure to a battery of 15 intense footshocks in a 90-min session) on single-outcome and sensory-specific PIT.

## General Methods

### Subjects

Long Evan rats were used in all the experiments in this study (N=89); group sizes, and sexes are outlined below in individual methods sections. Rats were pair and triplicate housed and maintained on a reverse light-dark schedule. Experimental procedures were conducted during the dark phase of the cycle. For all experiments rats were food restricted to between 85% and 90% of their *ad libitum* weights and maintained at this weight for the duration (though see experiment 2 for exception in a final test).

### Apparatus

Two distinct contexts housed in separate rooms were used for fear conditioning (Ctx A) and for appetitive training and testing (Ctx B). Chambers differed in dimensions, patterned backdrops, olfactory cues, floor textures, and other internal features. All chambers were standard Med Associates operant chambers housed in sound attenuating cabinets. Ctx A was 29.53 x 24.84 x 18.67 cm (LxWxH) with metal rod grid floors (19, 4.7mm rods), a horizontal zig zagged pattern back drop, and a single house light. For olfactory distinctiveness gauze pads infused with clover leaf essential oil (Crafter’s Choice™ Clove Leaf EO-Certified 100% Pure 1050) were placed into the scat collection trays at the base of the chambers. Ctx B was 29.53 x 23.5 x 27.31 cm with wire mesh inserts (19-Gauge Wire Mesh Fence; 0.5” mesh size) placed over metal rod grid floors and a backdrop with randomly distributed differently sized squares or circles. For olfactory distinctiveness, citrus cilantro fragrance oil infused gauze pads were placed in the scat trays at the start of each session (Crafter’s Choice™ Citrus Cilantro Fragrance Oil 548). Internal features of Ctx B chambers included two retractable levers flanking a recessed food magazine, a single speaker above the magazine, and two flat lights above the levers on the front wall of the chamber. On the opposing wall, a single hooded houselight was situated at the top center. Food hoppers and syringe pumps were externally housed and connected to the magazine for delivery of pellet and liquid reinforcement.

### Shock trauma treatment

Rats were placed in Ctx A, where they received 15, 1mA footshocks delivered through the metal rods of chamber’s grid floor. The session lasted 90 minutes and shocks were delivered on variable-time 360 seconds (VT 360”) schedule. Control rats were exposed to the same context for 90 minutes without any shocks. All sessions were video recorded for subsequent quantification of freezing behavior.

### Single-Outcome Pavlovian-to-Instrumental Transfer (Training and Testing)

All appetitive training was conducted in Ctx B. First, rats received magazine training to learn to collect pellet reinforcements (Bioserv Dustless Precision Pellets, 45mg; product# F0021) from the recessed magazine. Each session lasted 20 minutes, in which 20 pellets were delivered on a VT 60” schedule. The number of magazine sessions differed across experiments; see individual experimental timelines for more details. Rats then learned to press an active lever to earn pellets while responses on a simultaneously available inactive lever had no consequence (lever positions counterbalanced). Initially, rats were trained on a continuous reinforcement schedule (CRF) in which every press earned a single pellet until reaching the acquisition criterion of 50 pellets in a 40 minute single session. Next, rats were trained on variable interval (VI) schedules of reinforcement in 20 minute sessions where the schedule was thinned across sessions as follows: VI 10”, VI 30”, and VI 60” (2, 2, and 4 sessions, respectively). At the start of each instrumental session the houselights turned on with simultaneous insertion of the levers and the session terminated with the light turning off and retraction of the levers. After instrumental training, rats received eight 55-min Pavlovian discrimination sessions, in which one conditional stimulus (CS; 2-min 80dB white noise or 2 kHz tone, counterbalanced) was reinforced (CS+) and one CS was nonreinforced (CS-). On each CS+ trial, 4 pellets were delivered on a VT 30” schedule. The first trial began 2 minutes after the session start and subsequent trials were presented after a variable 5 minute intertrial interval (ITI). Each CS was presented 4 times in a quasi-random order. Each session began with the house light turning on and ended 2 minutes after the last trial with the house light turning off. Following training, rats were tested for Pavlovian-to-instrumental transfer in 41 minute sessions under extinction conditions. Each test began with the house light turning on and insertion of both levers, then 10 minutes into the session the first CS trial began, and subsequent trials were presented on a fixed 2-minute ITI. Each CS was presented 4 times in a quasi-random order and the session ended 1 minute after termination of the last trial. Rats were tested in 2 separate sessions on 2 consecutive days.

### Sensory-Specific Pavlovian to Instrumental Transfer (Training and Testing)

As with SO-PIT, all training and testing was conducted in Ctx B. Two unconditional stimulus outcomes were used for reinforcement, Bioserv Dustless Precision Pellets (45 mg) and 20% sucrose (0.1 ml). Magazine training was conducted separately for each outcome using the same parameters as described above. Next rats were trained to press one lever for pellets and one for sucrose in separate sessions. Instrumental training proceeded as described for SO PIT starting with CRF, ending with VI 60” (lever-outcome assignments counterbalanced). Next rats underwent Pavlovian conditioning to associate one CS with pellets and the other with sucrose; a white noise and flashing light (0.25” on/off) were used as distinct CSs (CS-outcome assignments counterbalanced). Each CS was presented for 2 minutes per trial during which 4 US deliveries were made on a VT 30” schedule. Trials were separated by variable 5-min ITIs. The first trial was presented 2 minutes into the session and the session ended 2 minutes following termination of the last trial. Each CS-O pair was trained in separate 27 minute sessions (8 sessions per CS-O pair). Rats were given instrumental reminder sessions for each R-O association one day before each test. Test sessions were identical to that described in SO-PIT. Rats were tested in 3 sessions with instrumental reminder sessions between each test. After session 3, rats were given instrumental reminder training on the following day, then food restriction was lifted and rats were tested one day later for a final *ad libitum* SS-PIT test.

### Data Analysis

For analysis of freezing behavior, videos were scored manually by visually sampling behavior for 2s every 10 seconds during the 3-min immediately prior to each shock delivery (or yoked time period for controls). One video in Experiment 2 containing 3 females was corrupted during the recording and therefore data from shock conditioning were lost for these subjects. For appetitive training and testing, data were collected via automated response measures recorded by the Med Associates software controlling the operant chamber machinery. For statistical analyses, ANOVAs, unpaired t-tests, and Holm-Sidak’s (HS) multiple comparisons were used planned and post-hoc comparisons and are specified in more detail in the results section.

## Experiment-Specific Methods

### Experiment 1: Effects of Massive Shock on SO-PIT in Males

Male Long Evans rats (N=32) approximately 70 days old at the start of the experiment were purchased from Charles River Laboratories and shipped to the OHSU animal facility one week before the start of the experiment. Rats were food restricted to 85-90% of their *ad libitum* bodyweights for the duration of the experiment. Rats were split into 4 groups of 8 rats. Pre-Ctrl and -Shock groups received either control treatment or massive shock (Ctx A), respectively, one day prior to the start of appetitive training (Ctx B). Post-Ctrl and Shock groups received either control treatment or massive shock (Ctx A), respectively, the day after the final Pavlovian conditioning session, one day prior to testing. Testing was conducted on the same day for all rats. Rats received 2 test sessions over 2 consecutive days. Appetitive training and testing for SO-PIT was conducted as described above.

### Experiment 2: Effects of Massive Shock on SS-PIT in Males and Females

Female (N=22) and male (N=35) Long Evans rats were bred at OHSU from breeders purchased from Charles River Laboratories. Rats were weaned at postnatal (PN) day 23. At PN 70 or older, rats were food restricted and the following day given either control treatment or massive shock in Ctx A. The following day appetitive training for SS-PIT began in Ctx B as described above.

## Results

### Experiment 1: Effect of a battery of footshocks on SO-PIT

#### Pre-training fear conditioning

Freezing behavior in the Pre-Shock group was evident by the second shock and reached asymptotic levels by the third shock, whereas freezing behavior was expectedly absent in the Pre-Ctrl group (Supplemental Fig 1a. Two-way RM ANOVA, main effect of treatment, F_(1, 14)_ = 361.1, p<0.01; main effect of time, F_(1, 196)_ = 21.41, p<0.01; time by treatment interaction, F_(14, 196)_ = 21.41, p<0.01).

**Figure 1.**
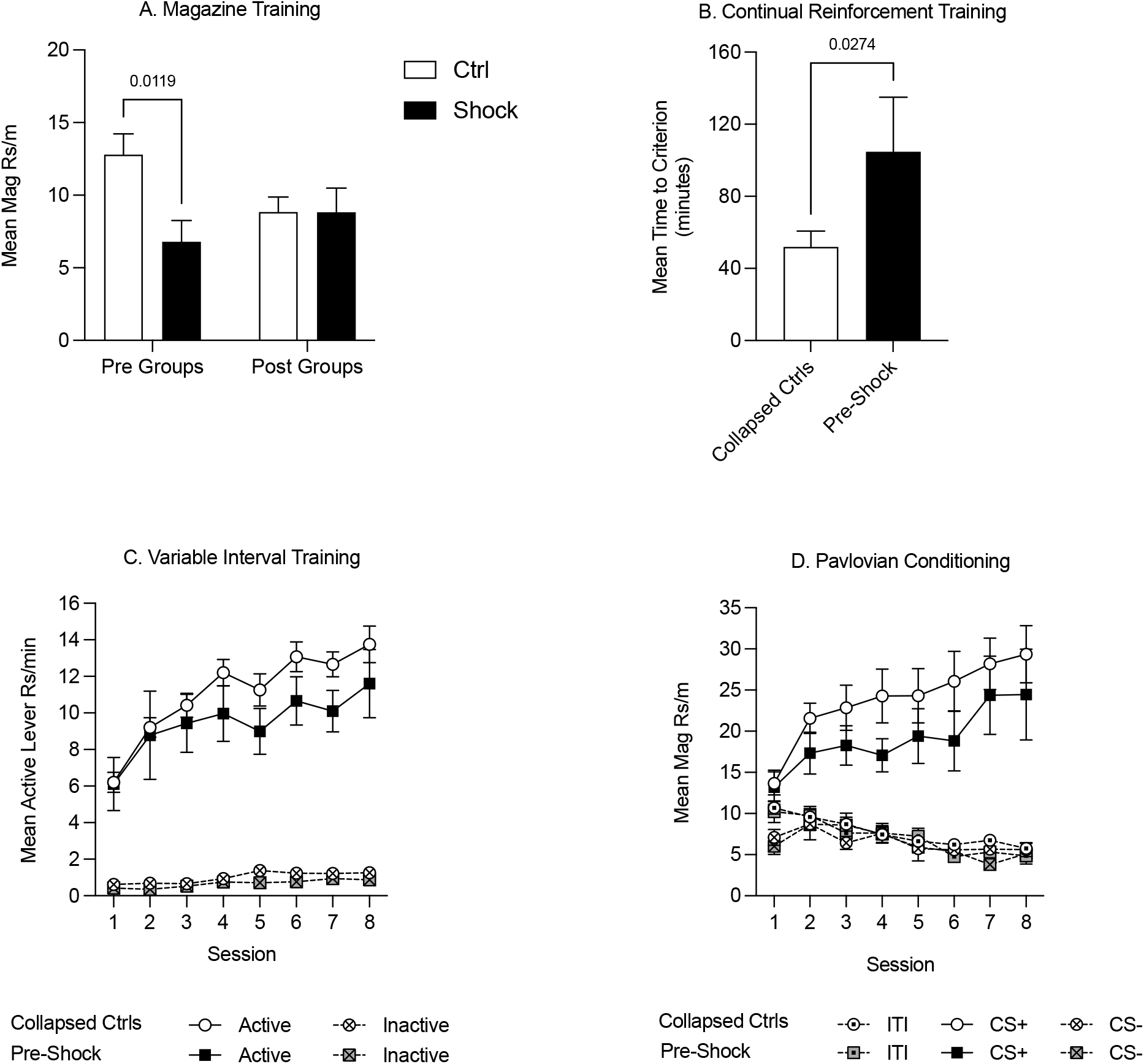
Appetitive training in Context B in Experiment 1. Data are collapsed between Pre-Ctrl, Post-Ctrl and Post-Shock groups when measures did not differ between these groups. A) Magazine training was suppressed in the pre-shock group relative to the pre-control group and there were no differences between the post groups. B) Acquisition on a continuous reinforcement schedule was slower in the pre-shock group compared to the other groups. C) During variable interval training, response rates increased across sessions and did not differ between Pre-Shock rats and the Collapsed Ctrl. Inactive lever responding was significantly lower than active lever responding across sessions. D) During Pavlovian conditioning, the mean rate of magazine responding during CS+ presentations increased across sessions and there were no group differences between Pre-Shock and Collapsed Ctrl Groups in responding.

#### Magazine, continuous reinforcement, and VI training

For all training data, when Pre-Ctrl, Post-Ctrl, and Post-Shock groups did not differ (p>0.05), these groups were collapsed (Collapsed Ctrl; i.e., all of the groups that had not received shock prior to training). Magazine training began the next day. As can be seen in Figure 1, rats that were shocked in Ctx A prior to appetitive training showed less magazine responding compared to the rats that received control context exposure in Ctx A (Figure 1a. Two-way RM ANOVA, significant treatment x time of treatment interaction, F_(1, 28)_ = 4.38, p<0.05; HS post-hoc comparison: Pre-Ctrl vs Pre-Shock, t_(28)_ = 2.98, p=0.01). During CRF training, the time to reach the acquisition criterion was delayed in the Pre-Shock group compared to the Collapsed Ctrl group (Fig 2b. t_(30)_ = 2.31, p=0.02). Active lever pressing increased across VI sessions in both groups and despite the ordinal difference in the groups there was no significant difference in active lever responding between the groups (Fig 1c. Two-way RM ANOVA, main effect of session, F_(7, 210)_ = 16.67, p<0.01; no effect of group, p=0.45; no session by group interaction, p=0.97). Inactive lever pressing was very low throughout training, but systematically increased without any group differences during the last 4 VI 60” sessions (Fig 1c. Two-way RM ANOVA, main effect of session, F_(7, 210)_ = 5.12, p<0.01; no effect of group, p=0.09; no session by group interaction, p=0.73). In both the Collapsed Ctrl and Pre-Shock groups responding on the active lever was significantly higher than the inactive lever and this difference increased over sessions (Fig 1c. Two-way RM ANOVA, main effect of session, Collapsed Ctrl, F_(7, 161)_ = 67.94, p<0.01; Pre-Shock, F_(7, 49)_ = 4.03, p<0.01; main effect of lever, Collapsed Ctrl, F_(1, 23)_ = 296.00, p<0.01; Pre-Shock, F_(1, 7)_ = 47.64, p<0.01; session by lever interaction: Collapsed Ctrl: F_(7, 161)_ = 19.84, p<0.01; Pre-Shock, F_(7, 49)_ = 2.71, p=0.02).

**Fig 2.**
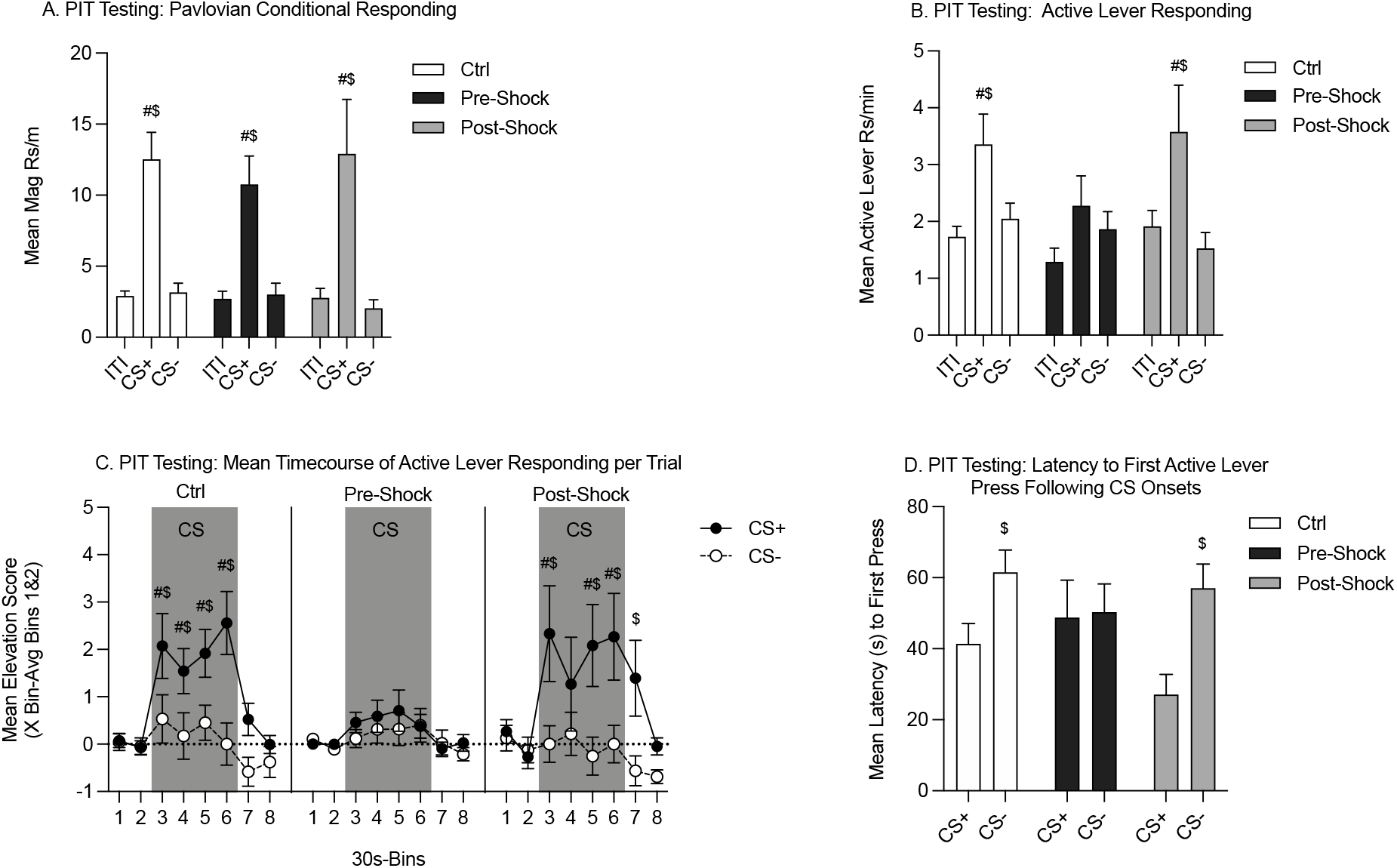
Single-outcome Pavlovian-to-instrumental transfer tests in Context B in Experiment 1. The Pre-Ctrl and Post-Ctrl groups did not differ and were therefore collapsed. Data are presented as means from Tests 1 and 2. A) Pavlovian conditional responding during PIT testing. All groups showed robust discrimination between CS+ and CS-. B) Active lever responding measuring the SO-PIT effect. In the Collapsed Ctrl and Post-Shock groups active lever responding was significantly greater during CS+ presentations than during the ITI and CS-. However, in the Pre-Shock the rate active lever responding did not differ between the CS+, ITI, and CS-periods. C) Mean time course of active lever from 60s pre-to 60s post-CS presentations. Data are presented as an elevation score (responding in the pre-CS period (bins 1 and 2) subtracted from each bin). In the Collapsed Ctrl and Post-Shock groups, active lever responding was greater during CS+ vs CS-presentations throughout the 2m CS window. In the Post-Shock group this effect carried over into the first 30s after CS offset. In contrast, the Pre-Shock group did not show any significant increase active lever responding during CS+ vs CS-presentations at any point within the CS window. D) Active lever response latency during PIT testing. In the Collapsed Ctrl and the Post-Shock groups, the latency between CS+ onset and the first active lever response was significantly shorter than the latency to press the active lever following CS-onset. In contrast, in the Pre-Shock group, there was no difference in the latency to first press following the CS+ vs the CS-onsets. ($: CS+ vs CS-, p<0.05; #: CS vs ITI, p<0.05)

#### Pavlovian Conditioning

During the eight sessions of appetitive Pavlovian conditioning, all groups responded more to the CS+ than to the CS- (Fig 1d. Two-way RM ANOVA, main effect of CS: Collapsed Ctrl: F_(2, 46)_ = 52.85, p<0.01; Pre-Shock: F_(2, 14)_ = 22.51, p<0.01; CS by session interaction: Collapsed Ctrl: F_(14,322)_ = 11.90, p<0.01; Pre Shock: F_(14, 98)_ = 4.59, p<0.01). As expected, magazine responding to the CS+ increased systematically across sessions and response rates between the Collapsed Ctrl and Pre-Shock Groups did not differ (Fig 1d. Two-way RM ANOVA, main effect of session, F_(7, 210)_ = 6.32, p<0.01; no effect of group, p=0.32; no session by group interaction, p=0.88). Magazine responding during the ITI and CS-did not differ between the Collapsed Ctrl and Pre-Shock groups and was expectedly low across training (Fig 1d. Twoway RM ANOVA, no effect of group: ITI: p=0.48; CS-: p=0.82). ITI responding significantly decreased across sessions, whereas CS-responding varied across days, but not systematically (Fig 1d. Two-way RM ANOVA, main effect of session: ITI: F_(7, 210)_ = 13.31, p<0.01; CS-: F_(7, 210)_ = 3.25, p<0.01).

#### Post-training fear conditioning

For the Post-Ctrl and Post-Shock groups, rats were given context exposure or received massive shock in Ctx A the day following the final appetitive Pavlovian conditioning session (the Pre-groups were left undisturbed in their home cages on this day). The Post-Ctrl and Post-Shock were counterbalanced to ensure even distribution of CS-O assignments and to match levels of appetitive Pavlovian conditional responding (data not shown: Two-way RM ANOVA, CS+ responding. no effect of group, p=0.82, no group x session interaction, p=0.99). During Ctx A training, as expected the Post-Ctrl group did not freeze significantly across the context exposure session, whereas the freezing in the Post-Shock group emerged rapidly and was significantly increased by the second pre-shock period compared to the first (Supplemental Fig 1b. Two-way RM ANOVA, main effect of group, F_(1, 14)_ = 204.2, p<0.01; main effect of time, F_(1, 196)_ = 7.88, p<0.01; time by group interaction, F_(14, 196)_ = 7.88, p<0.01; HS Post-hoc comparison: Bin 1 v 2: t_(196)_ = 4.45, p<0.01).

#### SO-PIT Testing

On the following two days all groups were tested for SO-PIT in Ctx B (one session per day). Although overall responding was lower during Test 2, there were no interactions with test session, so the two tests were combined for analysis. There also were no differences between Pre- and Post-Ctrl groups, so those data were combined for analysis (data not shown, two-way RM ANOVA, no effects of groups: Pavlovian conditional responding: p=0.74; active lever responding: p=0.53).

During testing, we evaluated magazine entries and lever pressing during the Pavlovian CSs. Each group showed greater magazine responding during the CS+ compared to the ITI (Fig 2a. HS planned comparison: Ctrl: t_(58)_ = 5.18, p<0.01; Pre-Shock: t_(58)_ = 3.07, p<0.01; Post-Shock: t_(58)_ = 3.86, p<0.01). Furthermore, CS+ magazine responding was significantly greater than CS-magazine responding (Fig 2a. HS planned comparison: Ctrl: t_(58)_ = 5.02, p<0.01; Pre-Shock: t_(58)_ = 2.95, p<0.01; Post-Shock: t_(58)_ = 4.14, p<0.01). Together, these data show that all groups retained the CS+/CS- discrimination, without notable group differences in these response rates.

Next, we examined active lever responding to evaluate PIT. Both the Ctrl and the Post-Shock groups showed SO-PIT with CS+ presentations evoking greater active lever responding compared to CS-presentations, which did not augment responding above ITI levels. This SO-PIT effect, however, was absent in the Pre-Shock group (Fig 2b. HS planned comparison CS+ vs CS-: Ctrl: t_(58)_ = 2.81, p=0.01; Pre-Shock: p=0.62; Post-Shock, t_(58)_ = 3.11, p=0.01). Furthermore, in the Ctrl and Post-Shock groups active lever responding was greater during CS+ vs ITI periods, but this augmentation by the CS+ was absent in the Pre-Shock group (Fig 2b. HS planned comparison: Ctrl, t_(58)_ = 3.49, p<0.01; Pre-Shock, p=0.36; Post-Shock, t_(58)_ = 2.53, p=0.03). We also evaluated how SO-PIT manifest across the CS presentation window by looking at the mean time course of responding in 30s bins from 60s before to 60s post CS presentations. These data are presented as elevation scores, which were calculated for each individual by subtracting the mean response during the pre-CS window (Bins 1 and 2) from the response rate in a given bin. In the Ctrl group, SO-PIT was highly temporally specific and was sustained and strengthened across the entire 2-min CS window (Fig 2c (left panel). Two-way RM ANOVA, main effect of time, F_(7, 105)_ = 9.15, p<0.01; main effect of CS, F_(1, 15)_ = 6.81, p=0.02; significant time by CS interaction, F_(7, 105)_ = 3.23, p<0.01). In the Pre-Shock group, SO-PIT was entirely absent across the CS window and did not change systematically over time (Fig 2c (middle panel). Two-way RM ANOVA, main effect of time, F_(7, 49)_ = 2.80, p=0.02; no effect of CS, p=0.60; no time by CS interaction, p=0.91). Although SO-PIT was evident across the session in the Post-Shock group when comparing CS+ to CS-, when pre-CS response rates were incorporated, PIT was not evident in every CS time bin and this was enough variability to weaken the overall main effect of CS (Fig 2c (right panel). Twoway RM ANOVA, main effect of time, F_(7, 49)_ = 3.81, p<0.01; main effect of CS, F_(1, 7)_ = 4.08, p=0.08; time by CS interaction, F_(7, 49)_ = 2.05, p=0.07).

As a final measure of PIT, we examined the latency to lever press following CS+ and CS-onsets. Consistent with the rate data, the Ctrl and Post-Shock groups showed a shorter latency to press the active lever following CS+ vs CS-presentations, whereas the Pre-Shock group did not show any difference in the latency to press following CS+ vs CS-presentations (Fig 2d. HS planned comparison: Ctrl, t_(29)_ = 2.52, p=0.04; Pre-Shock, p=0.90; Post-Shock, t_(29)_ = 2.65, p=0.03).

In Experiment 1, we found that a battery of footshocks in one context led to an impairment in SO-PIT in a different context in male rats. Although there were deficits in acquisition of magazine approach, there were no effects of shock on acquisition of Pavlovian or instrumental responding. In addition, during testing, which was conducted under extinction conditions, discriminatory magazine responding to CS+ vs CS-was intact in all groups. Similarly, instrumental response rates were similar across all groups under these extinction conditions. When shock followed appetitive training, there were no profound effects on PIT testing, but when shock preceded training PIT was absent. In Experiment 2, we evaluate the effects of pre-training shock on sensory-specific PIT in male and female rats.

### Experiment 2: Effect of a battery of footshocks on SS-PIT in male and female adults

#### Pre-training fear conditioning

Ctx A treatment was conducted the day before the start of appetitive conditioning. In the Shock groups freezing during the pre-shock periods emerged rapidly in both males and females without pronounced sex differences (Supplemental Fig 2. Two-way RM ANOVA, main effect of trial, F_(14, 406)_ = 27.04, p<0.01; no effect of sex, p=0.16, trial by sex interaction, F_(14, 406)_ = 1.64, p=0.07). To further explore early sex effects on freezing, we focused on freezing behavior in the first 4 pre-shock periods. Here we found that females froze significantly less in these early pre-shock periods compared to males (Two-way RM ANOVA, main effect of sex, F_(1, 29)_ = 5.50, p=0.03). Freezing did not emerge in Ctrl Female or Male rats (Supplemental Fig 2. No main effect of trial or sex, and no interaction, ps>0.99).

#### Magazine, CRF, and VI training

Appetitive training in Ctx B began with magazine training; data were analyzed as the mean rate of responding across all sessions. There was an overall main effect of shock treatment (F_(1,57)_ = 7.15, p <0.01) with no reliable main effect of sex or interaction, ps=0.19 (Fig 3a.). Planned comparisons to evaluate the effect of shock within each sex found no differences in magazine entries in females (p=0.40), but shocked males made fewer entries than did control males (Fig 3a. HS planned comparison, t_(57)_ = 3.33, p<0.01). Next rats went through CRF training and data were analyzed by taking the mean time to acquire both instrumental responses. The time to reach the acquisition criteria did not differ between treatment groups or sex (Fig 3b. HS planned comparison Ctrl vs Shock: females and males, ps> 0.20). In the next phase, rats were trained on VI reinforcement schedules for 8 sessions. Data were analyzed as the mean rate of responding across both levers. In both sexes, response rates increased across sessions, were similar between Ctrl and Shock groups, and this did not interact with session (Fig 3c. Two-way RM ANOVA, main effect of session: Females: F_(7, 140)_ = 2.61, p=0.01; Males: F_(7, 259)_ = 5.64, p<0.01; no effects of treatment groups, and no interactions, ps>0.41). We also compared responding between sexes for each treatment group. In Ctrl, responding did not differ between sexes, but in the Shock groups, response rates in Females were lower than in Males overall (Twoway RM ANOVA, main effects of sex: Ctrl: p=0.27; Shock: F_(1, 32)_ = 4.35, p<0.05).

**Figure 3.**
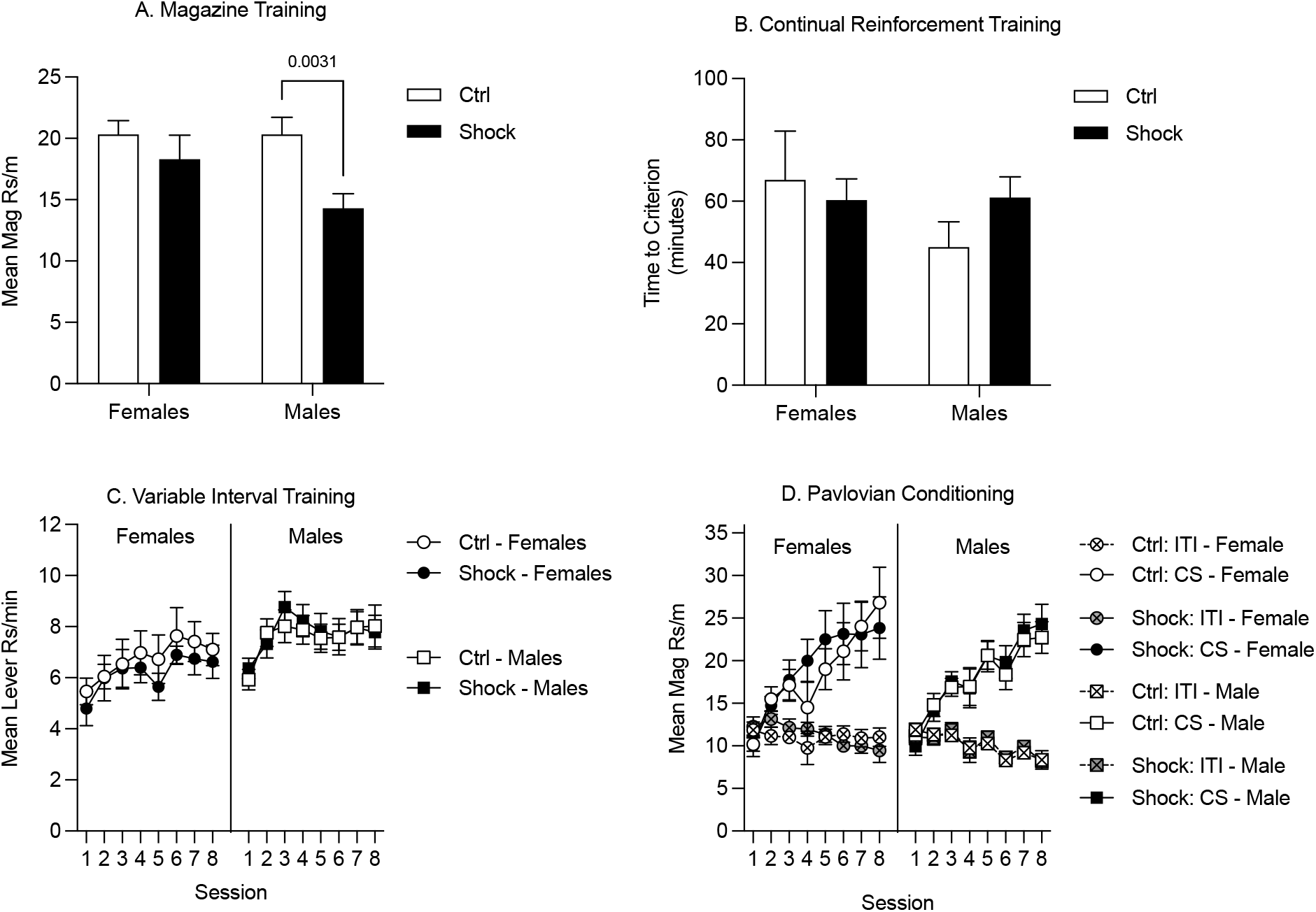
Appetitive training in Context B in Experiment 2. A) During magazine training, responding was similar between female Ctrl and Shock groups, whereas in males responding was suppressed in the Shock group relative to the Ctrl group. B) During CRF training, the time to reach the acquisition criteria did not differ between the Ctrl and Shock groups in either sex and there was no indication of a sex difference in acquisition times. C) During VI training, instrumental responding increased across sessions, but was not significantly different between the Ctrl and Shock groups in females or in males. (Though not shown directly here, responding in Shock groups was lower in females than males throughout training; there were no sex differences in responding in the Ctrl groups). E) During Pavlovian conditioning, magazine responding during CS+ presentations increased across sessions without any differences between the Ctrl and Shock groups and no differences or interactions with sex. Magazine responding during the ITI and CS-did not increase across session and there were not differences across groups.

#### Pavlovian Conditioning

In the last appetitive training phase rats underwent sensory-specific Pavlovian conditioning. These data were analyzed by averaging response rates across both CSs. In both sexes, magazine entries during the CSs increased across training sessions and did not differ between treatment groups (Fig 3d. Two-way RM ANOVA, main effect of session: Females: F_(7, 140)_ = 17.10, p<0.01; Males: F_(7, 259)_ = 29.46, p<0.01; no effects of treatment groups, Females, p=0.80; Males: p=0.87). In both sexes, response rates during the ITI were low and did not differ between treatment groups (Fig 3d. Two-way RM ANOVA, no effects of treatment group: Females, p=0.81; Males: p=0.94). Finally, we compared response rates between sexes for each treatment group and found no sex differences in either Ctrl or Shock groups (Two-way RM ANOVA, main effects of sex: Ctrl: p=0.83; Shock: p=0.71).

#### SS-PIT Testing

Rats were tested in 3 separate tests with instrumental reminder sessions on the days between testing. Initial analysis was conducted by averaging performance in these tests. Firstly, to evaluate the possibility of any notable sex differences or interactions between sex and trauma on the magnitude of SS-PIT, we calculated a mean score for SS-PIT magnitude by subtracting the average Diff lever response rate from the average Same lever rate. We found no effect of shock treatment, but a subtle sex difference with overall lower SS-PIT in females compared to males and this did not interact with shock treatment (Fig 4c. Two-way ANOVA, main effect of sex F_(1,57)_ = 3.36, p=0.07; no effect of treatment, p=0.48; no sex by treatment interaction, p=0.54).

**Figure 4.**
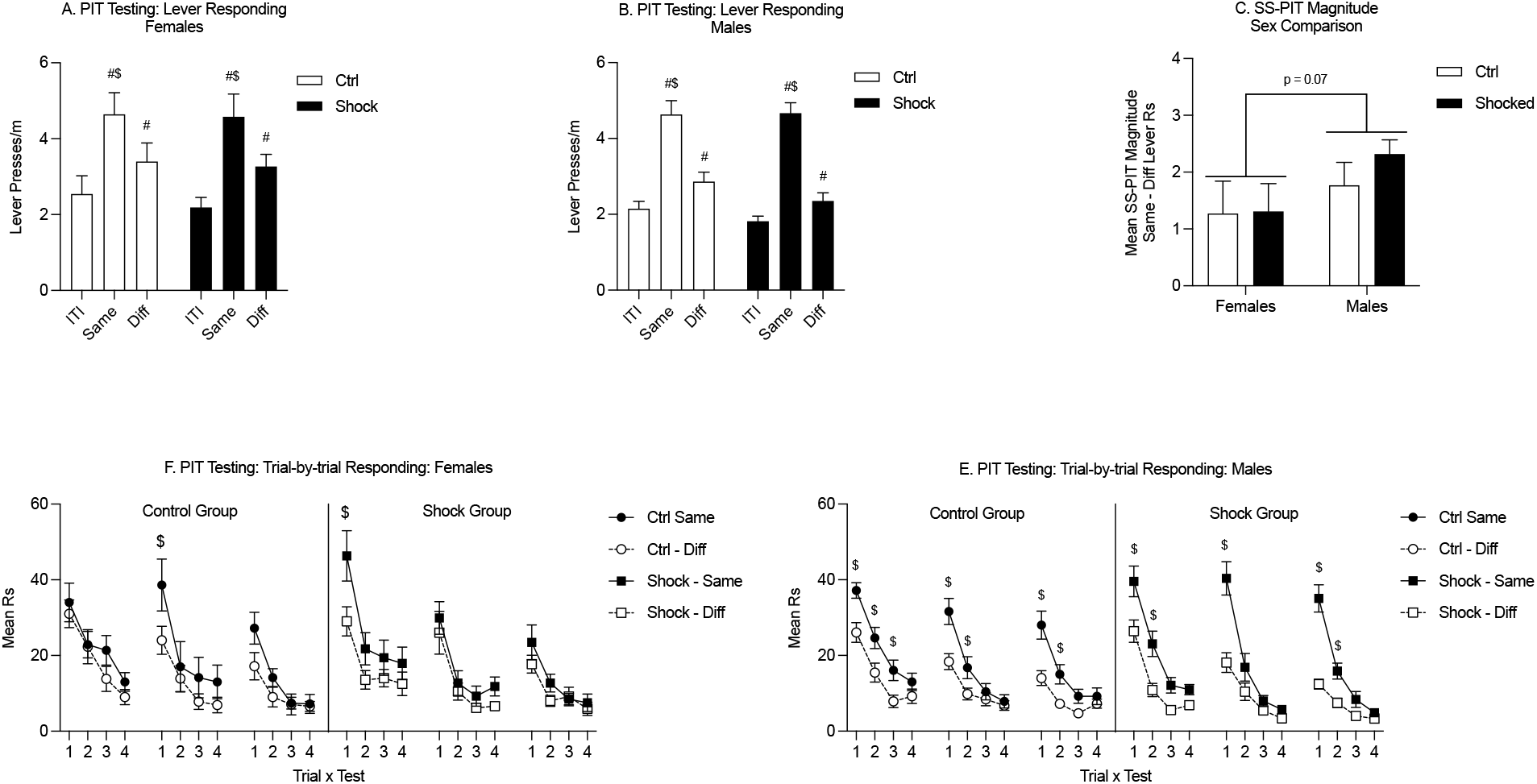
SS-PIT testing in Ctx B in Experiment 2. A) In females, SS-PIT was apparent in that lever responses on the Same lever were greater than responding on the Different lever in both Ctrl and Shock groups. General PIT was also evident in that Different lever responding was also significantly elevated above ITI lever response rates in both groups. B) In males, SS-PIT was also evident in both the Ctrl and Shock groups with Same lever responses being significantly higher than Different lever responses and as in the females, there was a notable General PIT effect with Different lever response rates being greater an ITI response rates. C) SS-PIT magnitudes did not did not differ between Ctrl and Shock groups, but overall there was a subtle indication of a sex difference, such that SS-PIT magnitudes were lower in females than males independent of shock treatment. D) In females the trial-by-trial SS-PIT effect was not strong and was only evident on 1/12 total trials across tests in Ctrl and Shock groups. Overall, response rates began high and extinguished across the trials within each test. E) In males in both the Ctrl and Shock groups the SS-PIT effect was apparent early in each test and disappeared with repeated trials. This pattern repeated across testing in a spontaneous recovery like effect. ($: Same vs Diff, p<0.05; #: significantly greater than ITI, p<0.05)

Next, to directly examine SS-PIT we looked at the means lever response rates across testing for ITI, Same lever, and different lever responding. In females both Ctrl and Shock groups exhibited sensory-specific PIT with a given CS preferentially increasing responding on the lever associated with the same outcome as the CS and there were no differences between the Ctrl and Shock groups (Fig 4a. Two-way RM ANOVA, main effect of lever response type (ITI, Same CS, Diff CS): F_(2,42)_ = 32.16, p<0.01; HS planned comparison: Same vs Diff: Ctrl, t_(42)_ = 3.07, p<0.01; Shock, t_(42)_ = 3.37, p<0.01; no effect of treatment, p=0.75). In addition to sensory-specific transfer (Same > Diff), both Ctrl and Shock females exhibited significant General transfer on the different lever such that CS presentations significantly elevated responding on the different lever over ITI response rates (Fig 4a. HS planned comparison: ITI vs Diff CS: Ctrl, t_(42)_ = 2.11, p=0.04; Shock, t_(42)_ = 2.79, p<0.01). The pattern of effect was the same in males, such that both Ctrl and Shock groups showed sensory specific transfer with greater same lever responding than different and no differences between Ctrl and Shock groups (Fig 4b. Two-way RM ANOVA, main effect of lever response type: F_(2,74)_ = 114.00, p<0.01; HS planned comparison: Same vs Diff: Ctrl, t_(74)_ = 6.37, p<0.01; Shock, t_(74)_ = 9.50, p<0.01; no effect of treatment, p=0.34). As in the females, both the Ctrl and Shock groups showed significant General transfer with presentation of CS evoking a significant increase in responses on the different lever from ITI responding (Fig 4b. HS planned comparison: ITI vs Diff CS: Ctrl, t_(74)_ = 2.58, p=0.01; Shock, t_(42)_ = 2.18, p=0.03). In summary Shock did not disrupt SS-PIT in either female or male rats.

To assess the expression of PIT across testing, we examined active lever responding during each test trial. In females, SS-PIT (Same>Diff) at the level of this trial-by-trial analysis was apparent in the Ctrl and the Shock groups (Fig 4e. Two-way RM ANOVA, main effect of CS: Ctrl: F_(1,9)_ = 5.08, p=0.05; Shock: F_(1, 11)_= 6.06, p=0.03). Sensory-specific PIT at a trial level was only evident in the female Ctrl group on the first trial of the second test, whereas in the female Shock group this SS-PIT by trial was only significant on the first trial of the first test. In contrast in males, the trial analysis revealed robust SS-PIT in both the Ctrl and Shock groups (Fig 4f. Two-way RM ANOVA, main effect of CS: Ctrl: F_(1, 16)_= 18.49, p<0.01; Shock: F_(1,20)_ = 50.53, p<0.01). In males, SS-PIT was strong at the beginning of each session, but extinguished over the course of test trials. This spontaneous recovery of SS-PIT effect occurred each day and the pattern of effects was parallel in the male Ctrl and Shock groups, with SS-PIT manifesting on 6/12 trials in the Ctrl group, and 5/12 trials in the Shock group. The trend toward an overall main effect of sex and these trial-by-trial data suggest a sex difference in the propensity of SS-PIT to manifest at a trial level independent of shock treatment, with males showing a more reliable trial-by-trial effect.

Given the subtle deficits in SS-PIT in females, we conducted an additional SS-PIT test, but under *ad libitum* conditions. The purpose of relieving food restriction for this test is based on the finding that SS-PIT is insensitive to alterations in satiation, whereas General PIT is weakened or entirely abolished by satiety (Balleine 1994, Corbit, Janak et al. 2007, Lingawi, Berman et al. 2022). Thus the goal here was to drive down any General PIT process that might be masking effects in females. During this final test, SS-PIT was absent in both Ctrl and Shock females, lever responding on the Same and Different levers to the CS in presentation did not differ in these groups (Fig 5a. Two-way RM ANOVA, main effect of lever response type (ITI, Same CS, Diff CS): F_(2,26)_ = 10.24, p<0.01; HS planned comparison: Same vs Diff: Ctrl, p=0.28; Shock, p=0.17; no effect of treatment, p=0.95). In contrast, in both male Ctrl and Shock groups SS-PIT was evident and with CS presentations evoking greater Same lever responses than Different lever responses (Fig 5b. Two-way RM ANOVA, main effect of lever response type (ITI, Same CS, Diff CS): F_(2,58)_ = 24.84, p<0.01; HS planned comparison: Same vs Diff: Ctrl, t_(52)_ = 2.82, p=0.01; Shock, t_(58)_ = 4.34, p<0.01; no effect of treatment, p=0.24).

**Figure 5.**
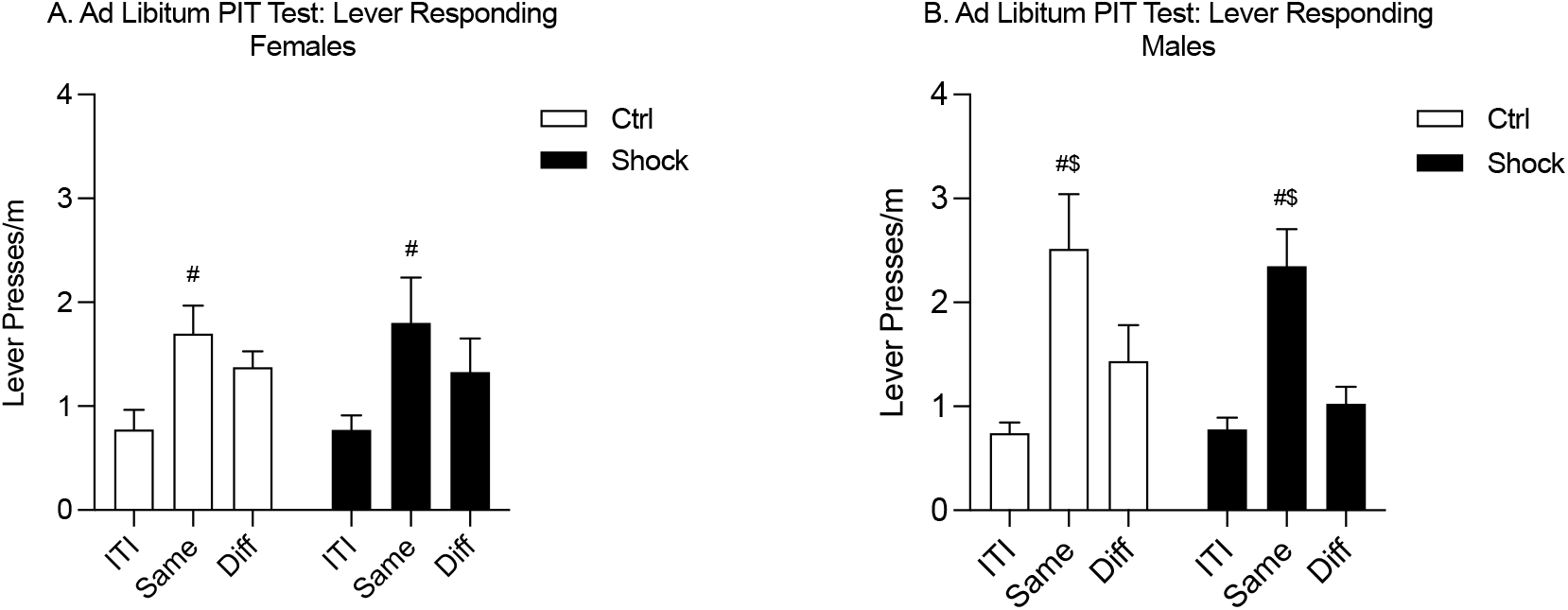
*<u>Ad Libitum</u>* SS-PIT test in Ctx B in Experiment 2. A) In females, the sensory-specificity of transfer was abolished in this ad libitum testing; Same and Different lever responses did not differ in either the Ctrl or Shock female groups. Same lever responses were greater than ITI in both groups. B) In males, SS-PIT was maintained under *ad libitum* test conditions and Same responding was significantly greater than Different lever responding. ($: Same vs Diff, p<0.05; #: significantly greater than ITI, p<0.05)

## Discussion

In this series of experiments we show that an acute intensely stressful experience prior to appetitive training persistently reduces single-outcome Pavlovian to instrumental transfer (SO-PIT) in adult males, but this same experience in male and female rats does not alter the expression of Sensory-Specific PIT (SS-PIT). These data show an acute highly stressful experience produces long lasting changes in general affective motivation for natural reward, as captured here with SO-PIT without overtly altering sensory-specific memory processes.

The effects of stress on learning, memory, and motivation are well documented in the literature. These studies have used acute stressors (such as footshock, yohimbine, or restraint stress) to evaluate the effects of stress on appetitive responding during or soon after a stressful experience (Mantsch, Baker et al. 2016). Typically, these acute stressors are thought to cause relatively short-term increases in stress. Other studies using chronic stressors (such as chronic variable stress, maternal separation, or long-term social isolation) have been shown to cause more persistent effects on consumption of and operant responding for natural rewards (Willner, Towell et al. 1987, Ghiglieri, Gambarana et al. 1997, Yan, Cao et al. 2010).

Our approach showing persistent changes of stress in appetitive responding is derived from the stress-enhanced fear learning (SEFL) procedure, in which a battery of footshocks in one context causes increased reactivity to a single footshock in a different context. SEFL occurs in contexts that have a long history of drug self-administration (PIzzimenti et al. 2017) and the battery of footshocks used to produce SEFL causes persistent changes in cue-induced reinstatement of methamphetamine-seeking (Pizzimenti, et al. 2017) and alcohol drinking (Meyer, et al. 2013).

These findings parallel two important aspects of PTSD as it often develops in humans: 1) a triggering traumatic event, and, 2) behavioral aberrations that persist long after the trauma. The results of work on drug-seeking in the context of SEFL parallel an important comorbidity seen in humans where PTSD is a major risk factor for substance use disorder. However, drugs of abuse alone trigger profound and persistent changes in appetitive motivation and the underlying circuitry, which makes it difficult to isolate the persistent impact of the acute trauma alone on appetitive processes. Furthermore, while many individuals suffering from PTSD experience substance use problems many do not, and understanding the impact of trauma on motivation for natural reward is important to understanding how PTSD manifests in such individuals and how to treat these comorbid symptoms.

In Experiment 1 we demonstrated that a in male rats a battery of footshocks before, but not after appetitive training blocks expression of SO-PIT. This effect was specific to the test phase; there were no differences in acquisition of the Pavlovian CS+ vs CS-discrimination or in acquisition of instrumental responding. During the PIT test, animals shocked prior to appetitive training did not show augmentation of instrumental responding during the CS+. The absence of SO-PIT was also apparent in the average time course of responding per trial and in the latency to press the levers following CS+ vs CS-presentations. Despite the failure for the excitatory properties of the CS+ to transfer to instrumental responding, these rats still showed reliable Pavlovian conditional responding, preferentially entering the food cup during CS+ vs CS-presentations. Thus, the failure for SO-PIT to manifest could not be explained by a failure to encode the Pavlovian discrimination. With all these considerations in mind, our data suggest that trauma can induce profound deficits in the ability for Pavlovian cues associated with natural reward to acquire general motivational properties in males.

Our finding here that shock prior to training induces deficits in SO-PIT are consistent with a study in humans showing that participants scoring high on anxiety and stress metrics did not show general PIT, which is thought to rely on the same mechanisms as SO-PIT, whereas those with low stress and anxiety did (Quail, Morris et al. 2017). Although our study is not examining anxiety and stress directly, the SEFL treatment itself is associated with long lasting elevations in stress and anxiety in rats and in humans a major feature of PTSD is abnormal elevations in stress and anxiety.

Although SO-PIT was disrupted by shock trauma administered prior to training, this same treatment after training, but before testing, had less impact on SO-PIT, with no effect overall but some subtle trial-by-trial deficits. This result is consistent with a study in rats showing that both single and multiple stressors administer either in the light or the dark phase of the circadian cycle did not block SO-PIT in Lister Hooded male rats (Pielock, Braun et al. 2013). It is notable that our procedure uses a very intense stress protocol which is distinct from that used in this prior study. Similarly, in humans, experimentally induced stress enhanced SO-PIT expression (Pool et al., 2015), thus it would seem that perhaps the immediate effects of stress on SO-PIT are variable and this warrants further investigation.

SO-PIT taps into basic motivational properties of Pavlovian cues, however it does not provide a clean read out of sensory-specific learning, memory, and motivation. In Experiment 1, all groups showed reliable conditional stimulus discrimination (CS+ vs CS-) regardless of shock history, arguing against trauma-induced problems with sensory memory encoding; however, the possibility remained that sensory-specific learning may be altered by a history of acute trauma. We therefore tested whether shock before training would alter the expression of SS-PIT, a clean measure of sensory-specific encoding. Here we used both female and male rats. Shock prior to training did not prevent the expression of SS-PIT in either male or female rats. Specifically, CS presentation selectively invigorated responding on the lever that previously shared the same outcome as the CS in presentation vs the lever that previously delivered a different outcome. This finding confirms that a history of trauma does not prevent the formation of distinct sensory-specific memories in either males or females. This finding is consistent with a previous study that used chronic unpredictable stress to examine the effects of stress on SS-PIT in male Wistar rats (Mordago et al., 2012). In the Mordago study, SS-PIT deficits were observed when testing was conducted shortly after the stress treatment period, but this deficit disappeared when rats were given a period of time off from stress treatment. In our design we used an acute stress and SS-PIT testing occurred more than 20 days after that stressful experience. Although we did not observe a short-term effect with SO-PIT in Experiment 1, it is possible that a test closer in time to the shock trauma would produce short-term deficits in SS-PIT.

In Experiment 2, there were several sex differences during the course of fear conditioning and PIT training and testing. First, during fear conditioning, females showed delayed acquisition relative to males early in the conditioning session. This delay may reflect early differences in expression of different behaviors (freezing in males, increased activity/escape in females; (Gruene, Flick et al. 2015). Despite some differences in rate of acquisition, asymptotic levels of freezing were identical in males and females.

Second, during magazine training in both experiments, which is the first appetitive training phase after shock, shocked males showed lower rates of magazine approach compared to control males, but in females the rates of magazine responding were similar in shock and control subjects. Unfortunately, we had no video record of the magazine training session, but we have found in other studies that males show more generalized freezing to this appetitive context than do females. This effect in Experiment 1 persisted into the first session of CRF training, but in Experiment 2, where there was more magazine training and thus more opportunity for generalized fear to extinguish prior to CRF training, there was no difference in CRF training. To the best of our knowledge, differences in freezing between males and females to a non-threatening novel context following SELF training has not been documented yet. During VI training, we did not see any sex differences between Ctrl groups, but in the Shock groups, females showed significantly lower rates of lever responding. This suggests that trauma may have differing effects on instrumental responding between sexes. It is possible that this effect is unique to responding under VI schedules or reinforcement or may reflect broader female specific trauma-induced deficits in instrumental responding. We did not see any indication of sex differences or interactions during Pavlovian conditioning.

Third, although females and males showed SS-PIT, the SS-PIT (i.e, the difference in Same vs Diff) was more consistent across testing in males, with females showing SS-PIT on only one of the 12 test trials. Moreover, when tested in a ad libitum state, females failed to show SS-PIT such that responding to the same vs different lever did not significantly differ. Given that SS-PIT has been shown to be insensitive to shifts in motivational states, particularly from hunger to satiety, we utilized this ad libitum test to determine to what extent females were relying on a sensory-specific process to solve the task (Balleine 1994, Corbit, Janak et al. 2007, Lingawi, Berman et al. 2022). The loss of sensory-specificity in transfer suggests that females, in our study, were relying more heavily on a generalized motivational process. It is possible these data represent a broad sex differences in reliance on or preferential engagement of sensory-specific vs general motivational information to guide cue triggered behavior. Together, these findings suggest that sex differences in fear conditioning, generalization, and SS-PIT are present, even if perhaps sometimes subtle. These effects warrant future study because differences in sex hormones may be enough to cause subtle changes in behavioral performance (Bangasser and Wicks 2017).

A noteworthy finding in Experiment 2 was within-session extinction and robust between-session spontaneous recovery of SS-PIT in males. On each test the SS-PIT (Same>Diff) effect was strongest in early trials growing weaker across trials and then re-emerging without any notable degradation upon retesting. Given that PIT testing is conducted under extinction conditions, the Pavlovian CSs are essentially given extinction training and while rats are retrained on the instrumental associations between tests, they never received additional CS-US pairing. Thus, the robust revival of the PIT effect on each subsequent test is quite similar to classic spontaneous recovery. The resistance of SS-PIT to Pavlovian extinction has previously been documented (Delamater 1996, Delamater, Schneider et al. 2017). However, the vast majority of SS-PIT studies that conduct multiple testing present the data in summary format potentially obscuring observation of this spontaneous recovery effect (Panayi and Killcross 2018, Alarcon and Delamater 2019, Lingawi, Berman et al. 2022, Sommer, Münster et al. 2022). To our knowledge, this is the first documentation of this effect.

Pizzimenti et. al. (2017) found that shock trauma prior to acquisition of methamphetamine self-administration resulted in enhanced cue-associated reinstatement of methamphetamine-seeking weeks after the initial trauma. These data are most parallel in design to our current study, but our current findings found a deficit rather than an enhancement in Pavlovian control of reward-seeking. The procedural differences between cue-associated reinstatement and PIT might explain some of the disparity between Pizzimenti’s finding and ours, however, there are alternative possibilities worth considering. Drugs of abuse exploit and engage the same circuitry that mediates appetitive motivation for natural rewards, but drugs of abuse themselves induce dramatic changes in plasticity and cues associated with drugs are more difficult to extinguish compared to cues associated with natural rewards (Jacobs, Hitchcock et al. 2022). Drugs of abuse also carry with them distinctively stressful properties. Thus, the effects observed by Pizzimenti may specifically reflect the unique plasticity induced by the interaction of acute trauma with drug abuse and drug withdrawal. Whereas in our study, the effects observed on PIT are more readily attributable to the acute stress alone, rather than stress or plasticity induced by the natural reward.

Another possibility is that our data reflect a stress-induced insensitivity to the motivational aspects of natural reward, which may render individuals more vulnerable to the motivational properties of drug reward. The suppressive effects of stress and depression on consumption and instrumental motivation for natural rewards has been well documented (Willner, Towell et al. 1987, Ghiglieri, Gambarana et al. 1997, Willner 2005, Yan, Cao et al. 2010). Thus our effects here on SO-PIT may represent an extension of these anhedonic effects to include the ability for Pavlovian CSs to acquire standard motivational properties. Given this possibility, perhaps acute trauma imparts a persistent anhedonia which we capture here in the form of blunted general affective motivation, which may render individuals more sensitive to the effects hedonic perturbation experienced by drugs resulting in a more salient experience thereby enhancing the incentive salience of drug associated stimuli. Ideally a direct comparison of drug versus natural reward using the same methodology (either PIT or cue-associated reinstatement) would be able to address his interesting discrepancy.

SO- and SS-PIT are dissociable both as psychological constructs and also rely on different brain circuits (Blundell, Hall et al. 2001, Corbit, Muir et al. 2001, Hall, Parkinson et al. 2001, Holland and Gallagher 2003, Corbit and Balleine 2005, Corbit, Janak et al. 2007, Corbit and Balleine 2011, Laurent, Leung et al. 2012, Cartoni, Balleine et al. 2016, Lingawi, Berman et al. 2022). Lesion and inactivation studies have shown that SO-PIT relies on the Nucleus Accumbens Core (NAc Core), and the Central Nucleus of the Amygdala (CeA) (Holland and Gallagher 2003, Corbit, Janak et al. 2007). In contrast, SS-PIT depends on the NAc Shell, and the Basolateral Amygdala (BLA), and the connection between these structures. Given this distinction at the neuronal level, it would seem likely that the persistent effects of trauma on SO-PIT, but not SS-PIT point to longterm trauma-induced plasticity in either the NAc Core and or the CeA. The absence of effects on SS-PIT does not rule out long term trauma induced changes in the NAc Shell and BLA, but suggest that whatever changes may be taking place they are not impacting the sub circuitry of these nuclei that are dedicated to the expression of this appetitive behavior (SS-PIT).

## Figure Captions

**Supplemental Fig 1.**
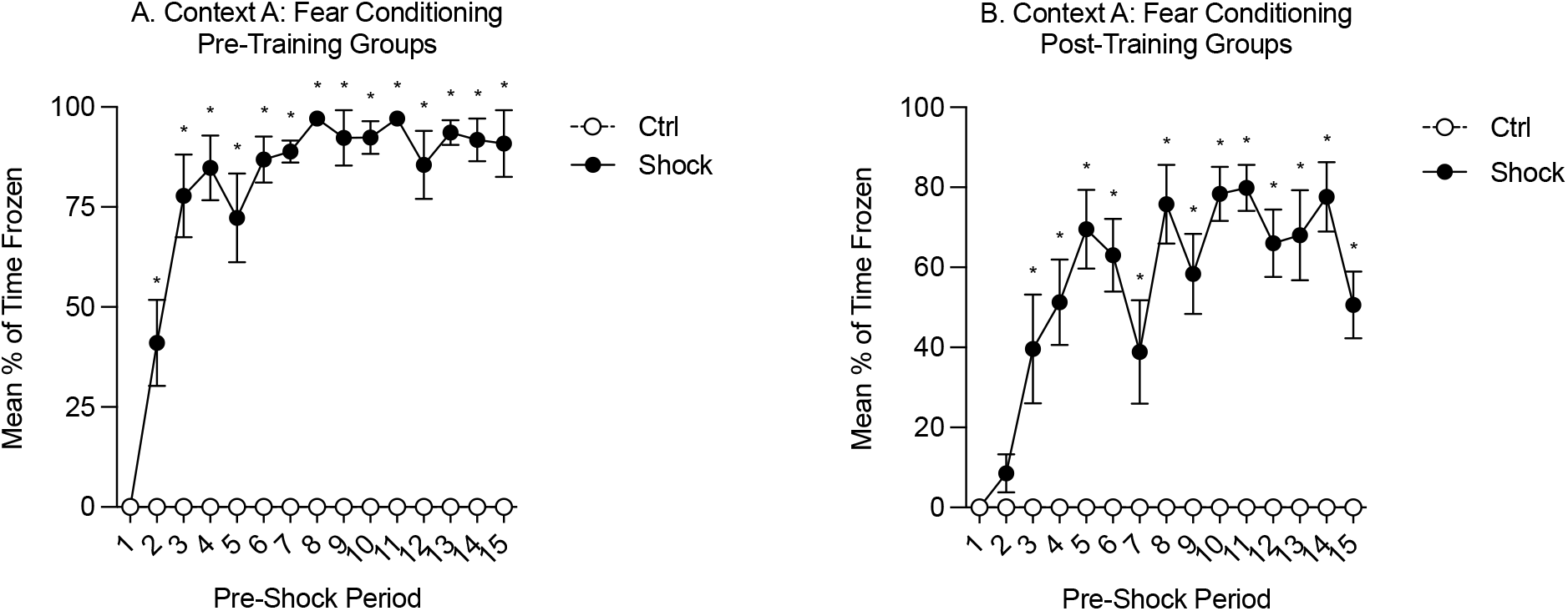
Freezing during fear conditioning in Context A in Experiment 1 when conditioning occurred (A) before or (B) after appetitive training. Freezing is shown during the 3-min periods before each of the 15 shocks (pre-shock period) or during yoked time periods for the control groups. *Ctrl vs Shock, p<0.001.

**Supplemental Fig 2.**
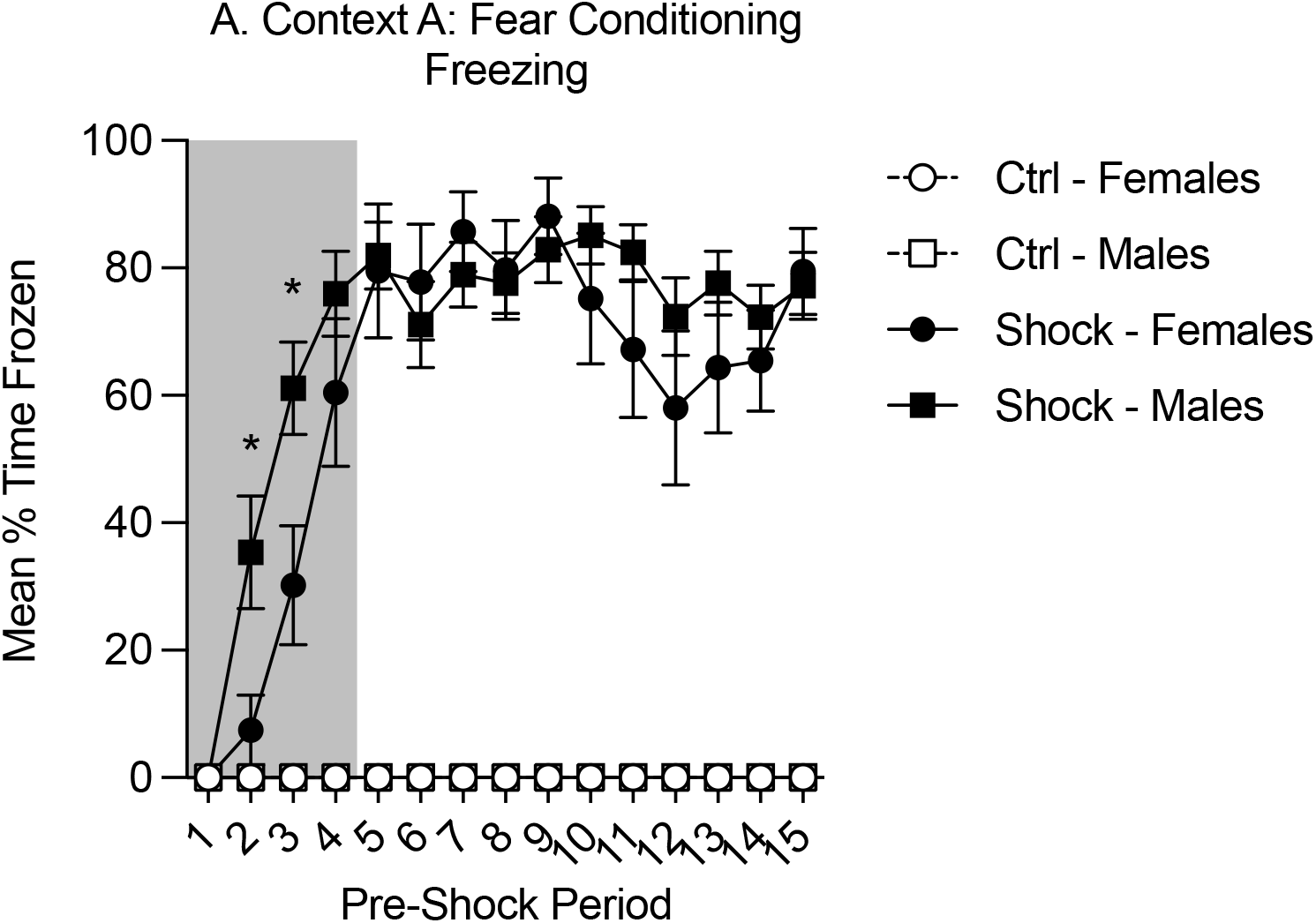
Freezing during fear conditioning in Context A in Experiment 2. Freezing is shown during the 3-min periods before each of the 15 shocks (pre-shock period) or during yoked time periods for the control groups. To examine early differences in freezing between sexes, we focused on the first 4 pre-shock periods. In the shocked groups, females showed lower rates of freezing in the pre-shock 2 and 3 windows. Overall shocked rats froze more than controls in both sexes (*: Female vs Male, p<0.01).

## Notes

Author Note: This work was supported by grants R01 DA047981 to KML and T32 DA007262 to RCD. Data and materials can be obtained from.

### Competing Interest Statement

The authors have declared no competing interest.

## References

Alarcon, D. E. and A. R. Delamater (2019). “Outcome-specific Pavlovian-to-instrumental transfer (PIT) with alcohol cues and its extinction.” Alcohol 76: 131–146.

Association, A. P. (2013). Diagnostic and statistical manual of mental disorders: DSM-5.

Balleine, B. (1994). “Asymmetrical interactions between thirst and hunger in Pavlovian-instrumental transfer.” Q J Exp Psychol B 47(2): 211–231.

Bangasser, D. A. and B. Wicks (2017). “Sex-specific mechanisms for responding to stress.” J Neurosci Res 95(1-2): 75–82.

Blundell, P., G. Hall and S. Killcross (2001). “Lesions of the basolateral amygdala disrupt selective aspects of reinforcer representation in rats.” J Neurosci 21(22): 9018–9026.

Brewerton, T. D. (2007). “Eating disorders, trauma, and comorbidity: focus on PTSD.” Eat Disord 15(4): 285–304.

Cartoni, E., B. Balleine and G. Baldassarre (2016). “Appetitive Pavlovian-instrumental Transfer: A review.” Neurosci Biobehav Rev 71: 829–848.

Chwastiak, L. A., R. A. Rosenheck and L. E. Kazis (2011). “Association of psychiatric illness and obesity, physical inactivity, and smoking among a national sample of veterans.” Psychosomatics 52(3): 230–236.

Corbit, L. H. and B. W. Balleine (2005). “Double dissociation of basolateral and central amygdala lesions on the general and outcome-specific forms of pavlovian-instrumental transfer.” J Neurosci 25(4): 962–970.

Corbit, L. H. and B. W. Balleine (2011). “The general and outcome-specific forms of Pavlovian-instrumental transfer are differentially mediated by the nucleus accumbens core and shell.” The Journal of neuroscience: the official journal of the Society for Neuroscience 31(33): 11786–11794.

Corbit, L. H., P. H. Janak and B. W. Balleine (2007). “General and outcome-specific forms of Pavlovian-instrumental transfer: the effect of shifts in motivational state and inactivation of the ventral tegmental area.” Eur J Neurosci 26(11): 3141–3149.

Corbit, L. H., J. L. Muir and B. W. Balleine (2001). “The role of the nucleus accumbens in instrumental conditioning: Evidence of a functional dissociation between accumbens core and shell.” J Neurosci 21(9): 3251–3260.

Coughlin, S. S., H. K. Kang and C. M. Mahan (2011). “Selected Health Conditions Among Overweight, Obese, and Non-Obese Veterans of the 1991 Gulf War: Results from a Survey Conducted in 2003-2005.” Open Epidemiol J 4: 140–146.

Delamater, A. (1996). “Effects of several extinction treatments upon the integrity of Pavlovian stimulus-outcome associations.” Animal Learning & Behavior 24(4): 13.

Delamater, A. R., K. Schneider and R. C. Derman (2017). “Extinction of Specific Stimulus-Outcome (S-O) Associations in Pavlovian Learning With an Extended CS Procedure.” J Exp Psychol Anim Learn Cogn 4(10).

Ghiglieri, O., C. Gambarana, S. Scheggi, A. Tagliamonte, P. Willner and M. G. D. Montis (1997). “Palatable food induces an appetitive behaviour in satiated rats which can be inhibited by chronic stress.” Behavioural Pharmacology 8(6).

Grubbs, J. B., H. Chapman and K. A. Shepherd (2019). “Post-traumatic stress and gambling related cognitions: Analyses in inpatient and online samples.” Addict Behav 89: 128–135.

Gruene, T. M., K. Flick, A. Stefano, S. D. Shea and R. M. Shansky (2015). “Sexually divergent expression of active and passive conditioned fear responses in rats.” eLife 4: e11352.

Hall, J., J. A. Parkinson, T. M. Connor, A. Dickinson and B. J. Everitt (2001). “Involvement of the central nucleus of the amygdala and nucleus accumbens core in mediating Pavlovian influences on instrumental behaviour.” Eur J Neurosci 13(10): 1984–1992.

Holland, P. C. and M. Gallagher (2003). “Double dissociation of the effects of lesions of basolateral and central amygdala on conditioned stimulus-potentiated feeding and Pavlovian-instrumental transfer.” Eur J Neurosci 17(8): 1680–1694.

Jacobs, D. S., L. N. Hitchcock, R. G. Williams and K. M. Lattal (2022). “Effects of a cue associated with cocaine or food reinforcers on extinction and postextinction return of behavior.” Behavioral Neuroscience 136(4): 307–317.

Kessler, R. C., W. T. Chiu, O. Demler, K. R. Merikangas and E. E. Walters (2005). “Prevalence, severity, and comorbidity of 12-month DSM-IV disorders in the National Comorbidity Survey Replication.” Archives of general psychiatry 62(6): 617–627.

Laurent, V., B. Leung, N. Maidment and B. W. Balleine (2012). “mu- and delta-opioid-related processes in the accumbens core and shell differentially mediate the influence of reward-guided and stimulus-guided decisions on choice.” J Neurosci 32(5): 1875–1883.

Lingawi, N. W., T. Berman, J. Bounds and V. Laurent (2022). “Sensory-Specific Satiety Dissociates General and Specific Pavlovian-Instrumental Transfer.” Front Behav Neurosci 16: 877720.

Mantsch, J. R., D. A. Baker, D. Funk, A. D. Lê and Y. Shaham (2016). “Stress-Induced Reinstatement of Drug Seeking: 20 Years of Progress.” Neuropsychopharmacology 41(1): 335–356.

Meyer, E. M., V. Long, M. S. Fanselow and I. Spigelman (2013). “Stress increases voluntary alcohol intake, but does not alter established drinking habits in a rat model of posttraumatic stress disorder.” Alcohol Clin Exp Res 37(4): 566–574.

Morgado, P., M. Silva, N. Sousa and J. J. Cerqueira (2012). “Stress Transiently Affects Pavlovian-to-Instrumental Transfer.” Front Neurosci 6: 93.

Panayi, M. C. and S. Killcross (2018). “Functional heterogeneity within the rodent lateral orbitofrontal cortex dissociates outcome devaluation and reversal learning deficits.” eLife 7: e37357.

Piazza, P. V., J. M. Deminiere, M. le Moal and H. Simon (1990). “Stress- and pharmacologically-induced behavioral sensitization increases vulnerability to acquisition of amphetamine self-administration.” Brain Research 514(1): 22–26.

Pielock, S. M., S. Braun and W. Hauber (2013). “The effects of acute stress on Pavlovian-instrumental transfer in rats.” Cognitive, Affective, & Behavioral Neuroscience 13(1): 174–185.

Pizzimenti, C. L., T. M. Navis and K. M. Lattal (2017). “Persistent effects of acute stress on fear and drug-seeking in a novel model of the comorbidity between post-traumatic stress disorder and addiction.” Learn Mem 24(9): 422–431.

Pool, E., T. Brosch, S. Delplanque and D. Sander (2015). “Stress increases cue-triggered “wanting” for sweet reward in humans.” J Exp Psychol Anim Learn Cogn 41(2): 128–136.

Quail, S. L., R. W. Morris and B. W. Balleine (2017). “Stress associated changes in Pavlovian-instrumental transfer in humans.” Q J Exp Psychol (Hove) 70(4): 675–685.

Rau, V., J. P. DeCola and M. S. Fanselow (2005). “Stress-induced enhancement of fear learning: An animal model of posttraumatic stress disorder.” Neuroscience & Biobehavioral Reviews 29(8): 1207–1223.

Rau, V. and M. S. Fanselow (2009). “Exposure to a stressor produces a long lasting enhancement of fear learning in rats.” Stress 12(2): 125–133.

Sommer, S., A. Münster, J.-A. Fehrentz and W. Hauber (2022). “Effects of Motivational Downshifts on Specific Pavlovian-Instrumental Transfer in Rats.” International Journal of Neuropsychopharmacology 25(3): 173–184.

Steins-Loeber, S., F. Lörsch, C. van der Velde, A. Müller, M. Brand, T. Duka and O. T. Wolf (2020). “Does acute stress influence the Pavlovian-to-instrumental transfer effect? Implications for substance use disorders.” Psychopharmacology (Berl) 237(8): 2305–2316.

Swinbourne, J. M. and S. W. Touyz (2007). “The co-morbidity of eating disorders and anxiety disorders: a review.” Eur Eat Disord Rev 15(4): 253–274.

Willner, P. (2005). “Chronic Mild Stress (CMS) Revisited: Consistency and Behavioural-Neurobiological Concordance in the Effects of CMS.” Neuropsychobiology 52(2): 90–110.

Willner, P., A. Towell, D. Sampson, S. Sophokleous and R. Muscat (1987). “Reduction of sucrose preference by chronic unpredictable mild stress, and its restoration by a tricyclic antidepressant.” Psychopharmacology (Berl) 93(3): 358–364.

Yan, H. C., X. Cao, M. Das, X. H. Zhu and T. M. Gao (2010). “Behavioral animal models of depression.” Neurosci Bull 26(4): 327–337.

